# MYC Serine 62 phosphorylation promotes its binding to DNA double strand breaks to facilitate repair and cell survival under genotoxic stress

**DOI:** 10.1101/2025.03.19.644227

**Authors:** Gabriel M. Cohn, Colin J. Daniel, Jennifer R. Eng, Xiao-Xin Sun, Carl Pelz, Koei Chin, Alexander Smith, Charles D. Lopez, Jonathan R. Brody, Mu-shui Dai, Rosalie C. Sears

## Abstract

Genomic instability is a hallmark of cancer, driving oncogenic mutations that enhance tumor aggressiveness and drug resistance. MYC, a master transcription factor that is deregulated in nearly all human tumors, paradoxically induces replication stress and associated DNA damage while also increasing expression of DNA repair factors and mediating resistance to DNA-damaging therapies. Emerging evidence supports a non-transcriptional role for MYC in preserving genomic integrity at sites of active transcription and protecting stalled replication forks under stress. Understanding how MYC’s genotoxic and genoprotective functions diverge may reveal new therapeutic strategies for MYC-driven cancers. Here, we identify a non-canonical role of MYC in DNA damage response (DDR) through its direct association with DNA breaks. We show that phosphorylation at serine 62 (pS62-MYC) is crucial for the efficient recruitment of MYC to damage sites, its interaction with repair factors BRCA1 and RAD51, and effective DNA repair to support cell survival under stress. Mass spectrometry analysis with MYC-BioID2 during replication stress reveals a shift in MYC’s interactome, maintaining DDR associations while losing transcriptional regulators. These findings establish pS62-MYC as a key regulator of genomic stability and a potential therapeutic target in cancers.

## Introduction

MYC is a master transcriptional regulator that impacts all cellular pathways involved in proliferation, differentiation, and response to cellular signals (Fernandez, Frank et al. 2003, Chen, Liu et al. 2018). Due to its participation in anabolic and stress-responsive biology, MYC deregulation is found in virtually all human cancers, is prognostic for patient survival, and is often responsible for chemotherapy resistance(Dang 2012, Gabay, Li et al. 2014, Stine, Walton et al. 2015, Kumari, Folk et al. 2017). To prevent MYC’s oncogenic effect in normal cells, MYC protein abundance and activity is tightly regulated with a half-life of about 15-30 minutes under physiological conditions(Hann and Eisenman 1984). MYC’s stability is primarily regulated through sequential phosphorylation events within MYC’s transactivation domain, Thr58 (pT58-MYC) and Ser62 (pS62-MYC) which impact its degradation through the ubiquitin-proteosome system(Sears, Nuckolls et al. 2000, Welcker, Orian et al. 2004, Sun, Li et al. 2021). Upon cell growth stimulation, MYC becomes transiently stabilized by RAS-induced and/or cyclin-dependent kinase-mediated phosphorylation at Ser62 resulting in increased MYC stability and activity(Lutterbach and Hann 1994, Farrell, Pelz et al. 2013). To trigger MYC’s degradation, subsequent phosphorylation at Thr58 by GSK3 or BRD4 initiates MYC’s engagement with the ubiquitin-proteosome system (Gregory, Qi et al. 2003, Devaiah, Mu et al. 2020).

Mechanistically, MYC canonically functions as a transcription factor that enables potent transcriptional amplification, intensifying the recruitment and assembly of multiple protein complexes at each stage of transcription(Das, Lewis et al. 2023). MYC-driven transcriptional amplification is accompanied by genomic burdens such as increased torsional stress, R-loop formation, Transcriptional-Replication Conflicts (TRCs), among others(Jha, Kouzine et al. 2023). To mitigate this increase in genomic stress, emerging and non-canonical functions of MYC have recently been described. Along with recruiting transcriptional machinery to active promoters, MYC nucleates a “topoisome” complex between topoisomerase 1 & 2 to relieve DNA torsional stress produced by the elevated transcription(Das, Kuzin et al. 2022). MYC has also been shown to facilitate the transfer of polymerase associated factor 1c (PAF1c) to stalled RNA polymerase, activating several chromatin modifying complexes and DNA repair to ensure high fidelity elongation(Endres, Solvie et al. 2021). In response to a variety of cellular stressors including transcriptional stress, replication stress, and proteolytic stress, MYC proteins have been described to multimerize and protect replication-fork stability, decrease R-loop formation, and terminate transcription, all to protect genomic stability in the presence of stress(Papadopoulos, Solvie et al. 2022, Solvie, Baluapuri et al. 2022). Accompanying these MYC multimers were DNA maintenance proteins which aligns with previous findings that the neuronal MYC paralog, MYCN, and MYC are capable of recruiting critical components of DNA repair such as BRCA1, the TRRAP-containing NuA4 complex, and the p400 helicase to active promoters(Frank, Parisi et al. 2003, Tworkowski, Chakraborty et al. 2008, Kim, Woo et al. 2010, Herold, Kalb et al. 2019, Papadopoulos, Solvie et al. 2022). Although these studies have highlighted emerging roles of MYC in safeguarding the genome under cellular stress, a direct role for MYC in mediating DNA repair has not been explored.

In this study, we investigated a direct role of MYC in DNA repair in cancer cells with a focus on pancreatic ductal adenocarcinoma (PDAC). Our previous research demonstrated that MYC pathway activity is high in a subset of patients with aggressive, liver metastatic PDAC, characterized by elevated replication stress and DNA repair signatures(Link, Eng et al. 2025). We performed further analyses and confirmed a strong correlation between MYC and the tumor’s response to genomic instability, both at the transcriptional and tissue levels. To investigate the cellular mechanisms behind this strong correlation, we employed a DNA double-strand break (DSBs)-specific proximity ligation assay to discover that MYC associates with DSBs and that genomic stress enhances MYC’s association with DSBs as well as repair proteins such as BRCA1 and RAD51. Furthermore, using a MYC-BioID2 proximity-dependent proteomic approach, we observed a shift in MYC’s interactome under replication stress, marked by a notable enrichment of DNA repair machinery. Mechanistically, we discovered that MYC’s association with DSBs is dependent on the phosphorylation at serine 62 (pS62-MYC). Blocking this phosphorylation with a phosphorylation-deficient MYC mutant (S62A-MYC) disrupts BRCA1 and RAD51 recruitment to DSBs, resulting in a reduction in DNA repair and overall cell survival. Together, our findings reveal a novel direct role for MYC in DNA damage repair, offering new insights into MYC’s involvement in genome maintenance that could be leveraged for new therapeutic strategies.

## Results

### MYC Activity Positively Correlates with Genomic stress, DNA repair and Poor Patient Survival in PDAC

Our recent study demonstrated that pancreatic ductal adenocarcinoma (PDAC) patients with tumors exhibiting higher molecular signatures of tolerance to replication stress are more likely to develop liver metastasis and experience poorer overall survival(Link, Eng et al. 2025). Furthermore, this study demonstrated that elevated MYC activity was associated with a tumor survival advantage under conditions of high replication stress and DNA damage. To explore this dataset of 218 primary tumors and 71 metastases further, we generated a replication stress gene set based on the intersection between known cell cycle, check point, and DNA replication genes. We found a significant positive correlation between this replication stress signature score and the hallmark MYC-V1 target pathway score (Figure 1A). Since tumor cells experiencing high proliferation and transcriptional activity are often deficient in biosynthetic activity, they face stalled and collapsed replication forks leading to DSBs and genomic damage (Vesela, Chroma et al. 2017, Dreyer, Upstill-Goddard et al. 2021). In agreement with the replication stress signature, elevated hallmark MYC-V1 targets pathway activity significantly positively correlated with the hallmark DNA repair pathway score in our patient PDAC tumor samples, suggestive of MYC’s emerging non-canonical function involved in maintaining genomic integrity (Figure 1B). We then stratified our PDAC tumor dataset into either high or low hallmark MYC-V1 pathway score cohorts and performed Virtual Inference of Protein-Activity Enrichment Regulon (VIPER)(Alvarez, Shen et al. 2016, Lachmann, Giorgi et al. 2016) analysis. Comparing common replication stress response and DNA damage repair regulons from the VIPER analysis revealed a robust enrichment in tumors with high MYC-V1 pathway scores compared to those with lower MYC-V1 pathway scores (Figure 1C). In PDAC, patient survival and efficacy of treatment is impacted, in part, by somatic alterations in DNA damage response (DDR) genes(Dreyer, Upstill-Goddard et al. 2021, Link, Eng et al. 2025). To investigate whether DDR alteration status affects MYC’s impact on patient survival, we separated our cohort into four categories that stratify based on high/low Hallmark MYC V1 target score and whether a patient has a detected somatic DDR alteration in their tumor. Patients with MYC-high tumors had poor survival regardless of tumor DDR status (Figure 1D, red vs blue lines); however, patients with DDR-altered MYC-low tumors have a significantly better survival probability over MYC-high DDR-altered tumors (Figure 1D). This suggests that high MYC activity promotes tolerance to the presence of DDR alterations, supporting aggressive tumors and poor patient outcome including resistance to DNA damaging chemotherapy, as 24% of our patients in our cohort received neoadjuvant and/or adjuvant chemotherapy(Link, Eng et al. 2025).

**Figure 1:**
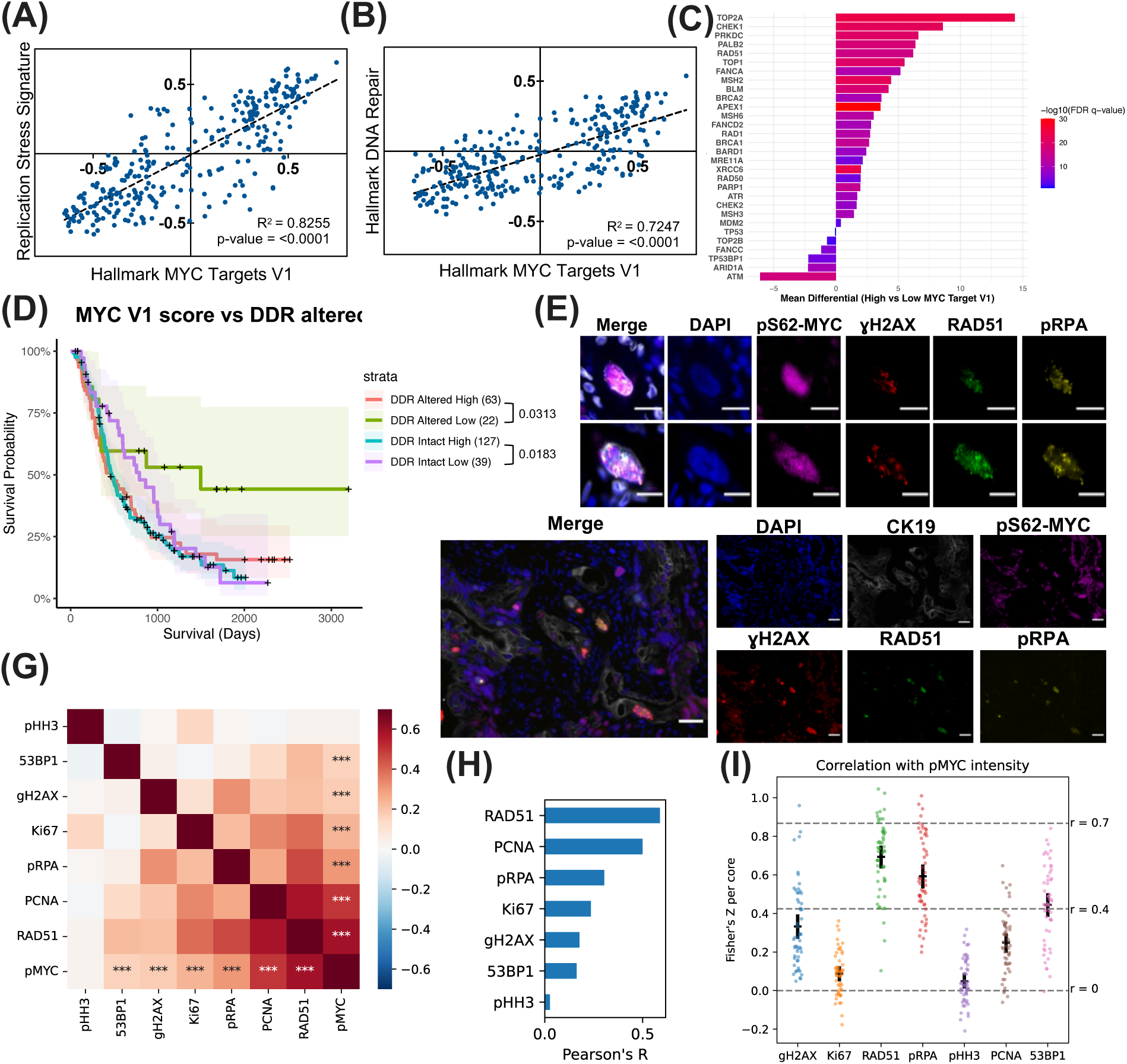
MYC Activity Positively Correlates with Genomic Damage and Poor Patient Survival in PDAC. (A) Pearson correlations of RNA expression of 289 primary and metastatic PDAC tumors comparing GSVA scores of hallmark MYC-V1 targets and a replication stress gene signature (B) as well as hallmark DNA repair pathway. (C) Mean differential VIPER regulon activity for DNA maintenance factors between tumors with high or low hallmark MYC-V1 target score, with coloring indicating FDR q-values from a one-way ANOVA. (D) K-M graph of overall survival for patients with high or low hallmark MYC-V1 target score stratified by tumors with (DDR altered) or without (DDR intact) known somatic alterations in DNA damage response-related genes. Shaded regions represent 95% CI. All P-values not shown are greater than 0.05. (E) Two close-up images of pS62-MYC positive and DNA damage marker positive cells. Scale bar, 26μm. (F) Representative images of cyclic immunofluorescence analysis of a single core from a PDAC tissue microarray. Scale bar, 32.5μm. (G) Pearson correlation (two-sided) of mean antibody intensities (***)FDR <0.001. (H) Ranked Pearson’s R correlation of mean antibody intensity. (I) Fisher’s Z transformed Pearson correlation r value per core of single cell mean intensity correlation with pS62-MYC intensity for all 54 cores across 34 patients. Error bars represent the 95% confidence interval. Dotted lines represent Pearson r value of 0, 0.4 and 0.7, respectively.

To investigate these findings at the protein level within human PDAC, we performed cyclic immunofluorescence (cycIF) on a patient-derived PDAC tissue microarray (TMA) with antibodies detecting markers of post-translationally active MYC (phospho-Ser62), cell proliferation, and DNA damage response. Consistently, we observed cytokeratin-19 positive tumor cells with overlapping staining for the activated pS62-MYC(Sears 2004, Hann 2006) and DNA damage markers RAD51, pRPA, and γH2AX (Figure 1E,F). When per cell intensities were quantified and ranked by correlation with pS62-MYC in tumor cells, in addition to the DNA damage proteins RAD51, pRPA, γH2AX and 53BP1, we observed significant correlation with cell cycle proteins PCNA and Ki67, suggestive of coupled proliferate and DNA repair (Figure 1G,H). However, when quantifying all 54 cores across 34 patients, the correlation between pS62-MYC and proliferation markers is not as strong while the correlation with DNA damage markers remain robust and significant (Figure 1I). Together, this data indicates that MYC expression correlates with markers for DNA damage response in PDAC tumors and impacts overall patient survival, particularly with patients with somatic DDR alterations.

### MYC is Detected in Proximity to DNA Double-Strand Breaks

To explore whether MYC plays a direct role in the molecular regulation of DNA breaks, we performed DNA Damage in situ Proximity Ligation Assay (DI-PLA) (Galbiati and d’Adda di Fagagna 2019), wherein a biotinylated DNA probe was ligated to DSBs, and proximity ligation assay (PLA) was conducted between biotin and MYC (Figure 2A). We treated an early passage patient-derived cancer cell line (ST-00024058), isolated from a resected PDAC tumor, with bleomycin and conducted DI-PLA. Bleomycin treatment resulted in a significant increase in DI-PLA puncta compared to DMSO control, indicating an enhanced proximity between MYC & DSBs in a patient-derived cancer cell line following DNA-damaging chemotherapy (Figure 2B). In a more precisely controlled system, we leveraged site-specific cleavage with a cas9 transfection with RNA guides targeting the 28S ribosomal DNA (rDNA) (van Sluis and McStay 2015). Targeting rDNA allows for signal amplification since there are approximately 400 copies of rDNA per cell. DI-PLA revealed a robust increase in association between MYC and cas9-induce DSBs compared to non-targeting control (NT) (Figure 2C). This suggests that MYC associates with DSBs generated by both chemotherapy and site-specific cleavage.

**Figure 2:**
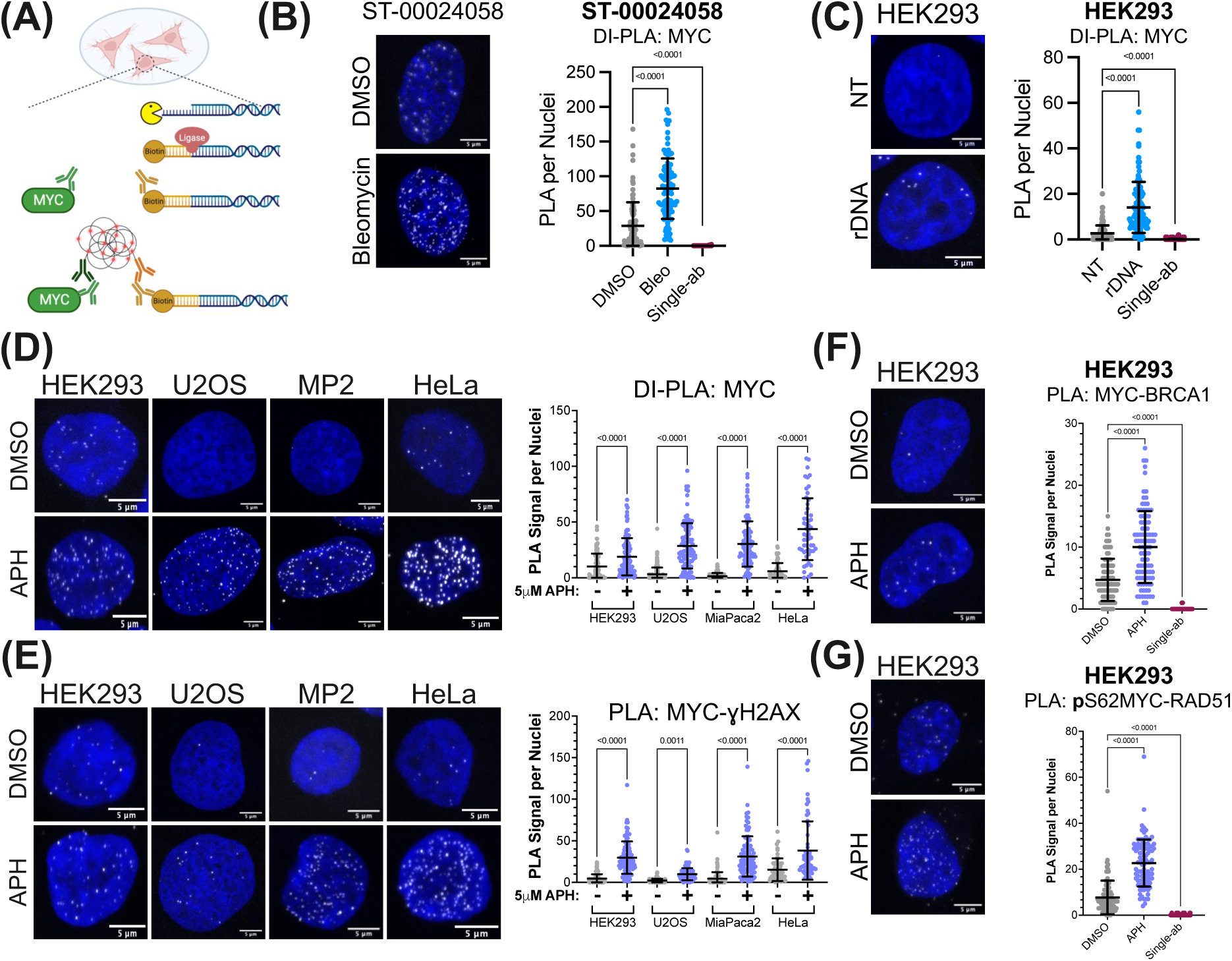
MYC is Detected in Proximity to DNA Double-Strand Breaks. (A) Schematic of DNA Damage in situ Proximity Ligation Assay (DI-PLA). (B) Left: Merged DI-PLA between MYC and biotin in a patient-derived PDAC cell line treated for 1 hour with 100 μg/mL bleomycin or DMSO. PLA puncta pseudo-colored in white. Right: quantification from three biological replicates of DI-PLA. Single antibody control combines PLA counts for both primary antibodies alone treated with bleomycin. The error bars show mean ±s.d. Statistical significance was determined by unpaired two-tailed t-test. (C) Left: Merged DI-PLA between MYC and biotin in HEK293 cells transfected with Cas9 protein and guide RNAs (sgRNA) targeting the 28S rDNA or non-targeting control (NT) for 8 hours. Right: quantification of DI-PLA. (D)Left: Representative images of DI-PLA between MYC and biotin in HEK293, U2OS, MIA PaCa-2, and HeLa cells treated with either 5μM APH or DMSO for 5-hours. PLA signal pseudo-colored in white. Left: Quantification of three biological replicates. (E) Left: Representative images of PLA between MYC and γH2AX. Right: Quantification of MYC and γH2AX PLA. (F) Left: Merged representative images of PLA between MYC and BRCA1 in HEK293 cells treated either DMSO or 5μM APH for 5-hours. Right: quantification of PLA. Single antibody control combines PLA counts for both primary antibodies alone treated with 5μM APH. (G) Left: PLA between pS62-MYC and RAD51 in HEK293 cells treated either DMSO or 5μM APH for 5-hours. Right: quantification of PLA. Note, MYC or pS62-MYC antibodies were chosen for PLA with RAD51 & BRCA1 antibodies based off cross-species compatibility.

As replication stress is a driver of DSBs and MYC expression correlates with pRPA expression, a marker of stalled replication forks (Figure 1D-G), we sought to determine whether MYC localization to DSBs is increased under high levels of replication stress in cells. To induce replication stress, cells were treated with aphidicolin (APH), an inhibitor of DNA polymerases α, δ, and ε, leading to the formation of vulnerable regions of single-stranded DNA, replication fork collapse, and the subsequent generation of DSBs(Vesela, Chroma et al. 2017). Consistent with direct DSB generation, APH treatment for 5-hours in HEK293, U2OS, MIA PaCa-2, and HeLa cells revealed a significant and robust increase in MYC’s association with replication stress-induced DSBs (Figure 2D). In agreement, the MYC-γH2AX PLA signal also increased following APH treatment in these cell lines, together highlighting the presence of MYC at DSBs following replication stress across multiple conventional cell lines (Figure 2E). To begin to understand the influence of MYC’s localization to DSBs on DNA repair, we tested whether APH induced an enrichment between MYC and known DNA repair proteins. BRCA1 binds DSBs and promotes homologous recombination directed repair and has been shown to bind to MYC, however, the mechanistic impact of this interaction remains to be clarified (Wang, Zhang et al. 1998, Li, Lee et al. 2002, Epasto, Pötzl et al. 2024). We observed a robust association between MYC and BRCA1 following 5-hour APH treatment, suggesting that this interaction is responsive to increased genomic damage (Figure 2F). Furthermore, we detected an APH-induced increase in association between the post-transcriptionally stable form of MYC (pS62-MYC) and RAD51 (Figure 2G); an interaction not previously reported in the literature. Taken together, our data suggests that MYC has a conserved capability of associating with DSBs and is able to interact with DNA repair proteins in response to DNA damage.

### MYC’s Interactome is Enriched for DNA Repair Proteins Under Replication Stress

To survey a broader and more unbiased enrichment of MYC’s interactions under replication stress, we generated stably expressed, doxycycline-inducible MYC-BioID2 in HEK293 cells for proximity-dependent labeling (Figure 3A). We treated these cells with either DMSO or APH and detected a total of 1,648 targets (Figure 3B). Our aim was to detect shifts in MYC’s interactome following 24-hour APH treatment. However, there were no statistically significant interactors enriched in APH treatment versus DMSO treatment following false discovery rate (FDR) correction (Figure 3C). This limited differential is likely attributed to a technical limitation of the BioID2 system, which requires an 18-hour incubation for biotinylation of proteins, whereas DNA damage response occurs on a much shorter timescale. Nevertheless, after background subtraction of the BioID2-only control and applying a ³2-fold differential abundance threshold and p=0.25 a shift in MYC protein target interactions in response to APH treatment is observed (Figure 3D). To better understand the functional differences of MYC interactors between DMSO and APH conditions, we conducted Gene Ontology (GO) enrichment using the BinGO tool in Cytoscape (Shannon, Markiel et al. 2003, Maere, Heymans et al. 2005, Doncheva, Morris et al. 2019). The interconnections of the enriched gene ontologies for MYC interactors under DMSO emphasize mechanisms which align with MYC’s canonical functions involved in gene expression, chromatin remodeling, and metabolism (Figure 3E). The APH-enriched MYC interactor ontologies highlight proteins involved in response to genomic damage, cell cycle regulation, and DNA repair (Figure 3F). These findings are consistent with recent studies demonstrating that under stress conditions, MYC shifts away from driving transcriptional processes to promoting DNA stability(Papadopoulos, Solvie et al. 2022, Solvie, Baluapuri et al. 2022). In agreement, when GO terms are ranked by adjusted p-value, we see the most significant terms for DMSO include processes involved in nucleotide metabolism and gene regulation while top terms for APH-induced MYC interactions pertain to genomic maintenance (Figure 3G). This data indicates that MYC’s interactome becomes more enriched for DNA damage response proteins under replication stress.

**Figure 3:**
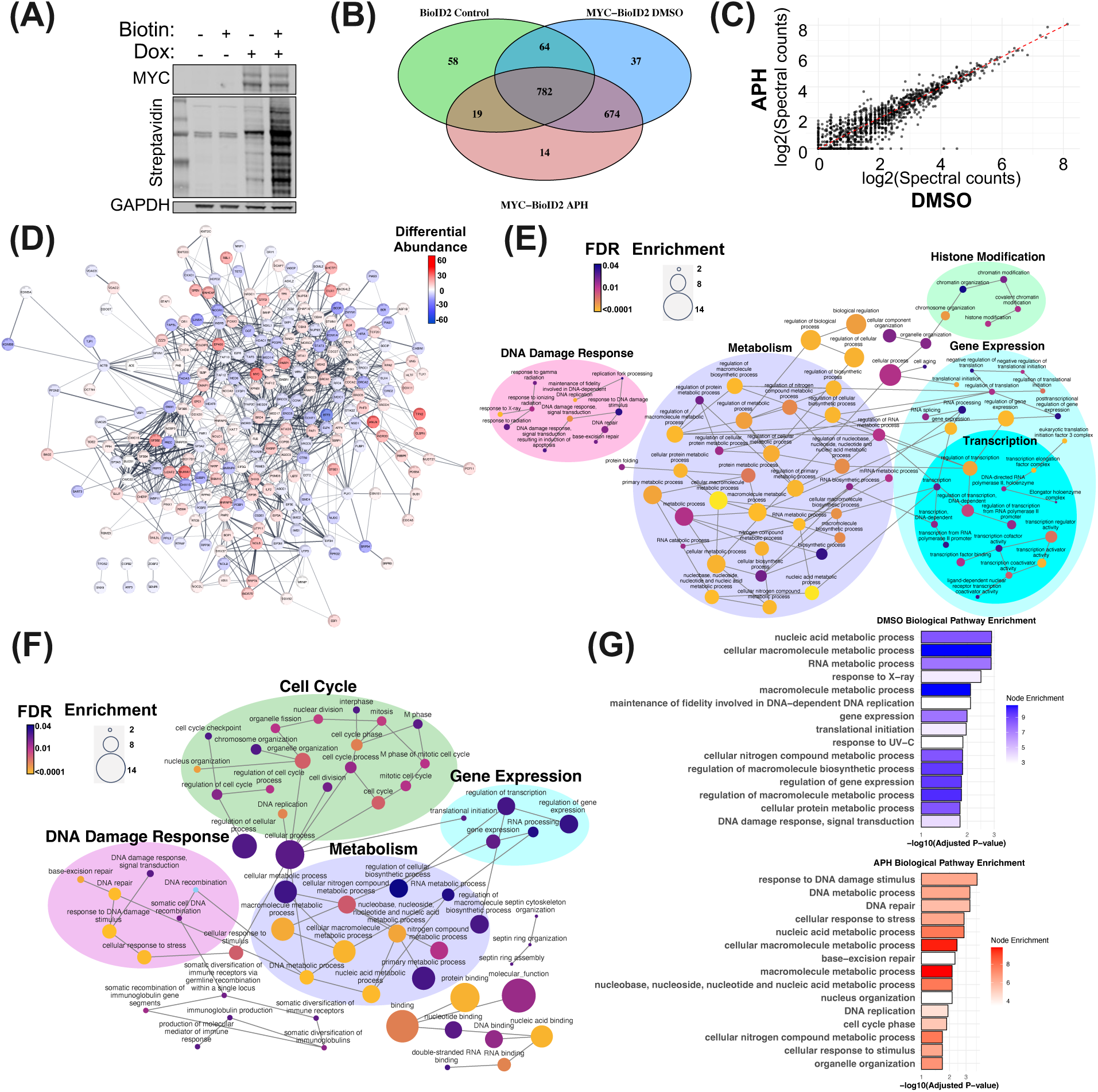
MYC Interactome is Enriched for DNA Repair Proteins Under Replication Stress. (A) Western Blot validation of inducible MYC-BioID2 construct in HEK293 cells treated with 1 μg/mL doxycycline for 24-hours and 50 μM biotin for 18-hours. (B) Venn diagram of biotinylated proteins detected in the different conditions, BioID2-only control (n=3), MYC-BioID2 treated with 24-hours of DMSO (n=3) or 5μM APH (n=2). (C) Scatter plot showing mean log2 value for the DMSO and APH conditions following background subtraction of BioID2-only. (D) Cytoscape interaction network analysis of MYC-BioID2 targets that pass a threshold of ≥2-fold differential abundance. Nodes colored based off mean spectral count differential abundance between APH (Red) and DMSO (Blue). BiNGO analysis showing the Gene Ontology (GO) enrichment network of DMSO (E) and APH (F). (G) Top 15 significantly enriched GO Biological pathway terms for DMSO (top) and APH (bottom).

### Serine 62 Phosphorylation of MYC Promotes its Association with DSBs

To begin to investigate whether post-translational modification of MYC could play a role in the functional switch in MYC activity upon genomic insult and its localization to DSBs and interaction with repair proteins, we investigated the impact of MYC’s phosphorylation status on this mechanism. Phosphorylation of MYC, particularly at threonine 58 and serine 62, have been shown to regulate its stability, target gene promoter binding, and spatial localization within the nucleus(Su, Pelz et al. 2018, Cohn, Liefwalker et al. 2020). In response to growth signals, MYC is transiently stabilized by the phosphorylation of serine 62, enhancing its engagement with target genes, including those poised at the nuclear periphery(Yeh, Cunningham et al. 2004, Vervoorts, Lüscher-Firzlaff et al. 2006, Farrell, Pelz et al. 2013, Su, Pelz et al. 2018). Processive phosphorylation at threonine 58 (pT58-MYC) destabilizes MYC, initiating its degradation via the proteosome(Gregory and Hann 2000, Welcker, Orian et al. 2004). To examine whether these phosphorylation sites in MYC could also affect its association with DSBs, we used doxycycline-inducible HEK293 cells which express hemagglutinin (HA) tagged wild-type MYC (WT-MYC), serine-to-alanine mutant MYC (S62A-MYC), or the threonine-to-alanine mutant MYC (T58A-MYC). Given the sequential nature of phosphorylation at these sites, with S62 phosphorylation preceding the phosphorylation of T58 by the processive GSK3 kinase (Gregory, Qi et al. 2003), S62A-MYC lacks phosphorylation at both sites, whereas T58A-MYC exhibits robust phosphorylation at S62 but lacks phosphorylation at T58 (Figure 4A). We performed HA-tagged DI-PLA following cas9-directed cleavage of the 28S rDNA. Consistent with endogenous MYC, we observed a robust increase in the association of ectopic WT-MYC and cas9-induced DSBs (Figure 4B). In contrast, S62A-MYC showed no statistically significant difference between non-targeting (NT) control and cas9-induced DSBs, while T58A-MYC exhibited a similar or even greater induction of MYC associated with DSBs compared to WT. Similar induction patterns were observed in a PLA between HA-tag and γH2AX (Figure 4C), demonstrating that serine 62 phosphorylation is a key determinant of MYC’s association with cas9-induced DSBs. To confirm these findings under replication-stress induced DSBs, we treated the MYC mutant cells with APH. WT-MYC showed a strong increased association with APH-induced DSBs compared to the DMSO control (Figure 4D). Likewise, T58A-MYC demonstrated a robust increase in association with APH-induced DSBs. While the S62A-MYC mutant also showed some increased association with DSBs, this was significantly less than WT-MYC and T58A-MYC. T58A-MYC retains persistent serine 62 phosphorylation due to resistance to PP2A-mediated dephosphorylation, unlike WT-MYC (Yeh, Cunningham et al. 2004, Arnold and Sears 2006). Notably, T58A-MYC displayed elevated association with DSBs even prior to APH treatment. Collectively, these data indicate that serine 62 phosphorylation plays an important role in MYC’s efficient association with DSBs in response to cas9- and APH treatment-induced DSBs. To verify that the difference in association with DSBs was not due to differences in expression levels, whole cell lysates of the three cell lines showed equal expression of the HA-tag MYCs (Figure 4E). In agreement with past findings of negative autoregulation (Kaur and Cole 2013), the lower molecular weight band of endogenous MYC was nearly undetectable when ectopic MYC is expressed, supporting that endogenous MYC did not compensate for the phosphorylation mutants.

**Figure 4:**
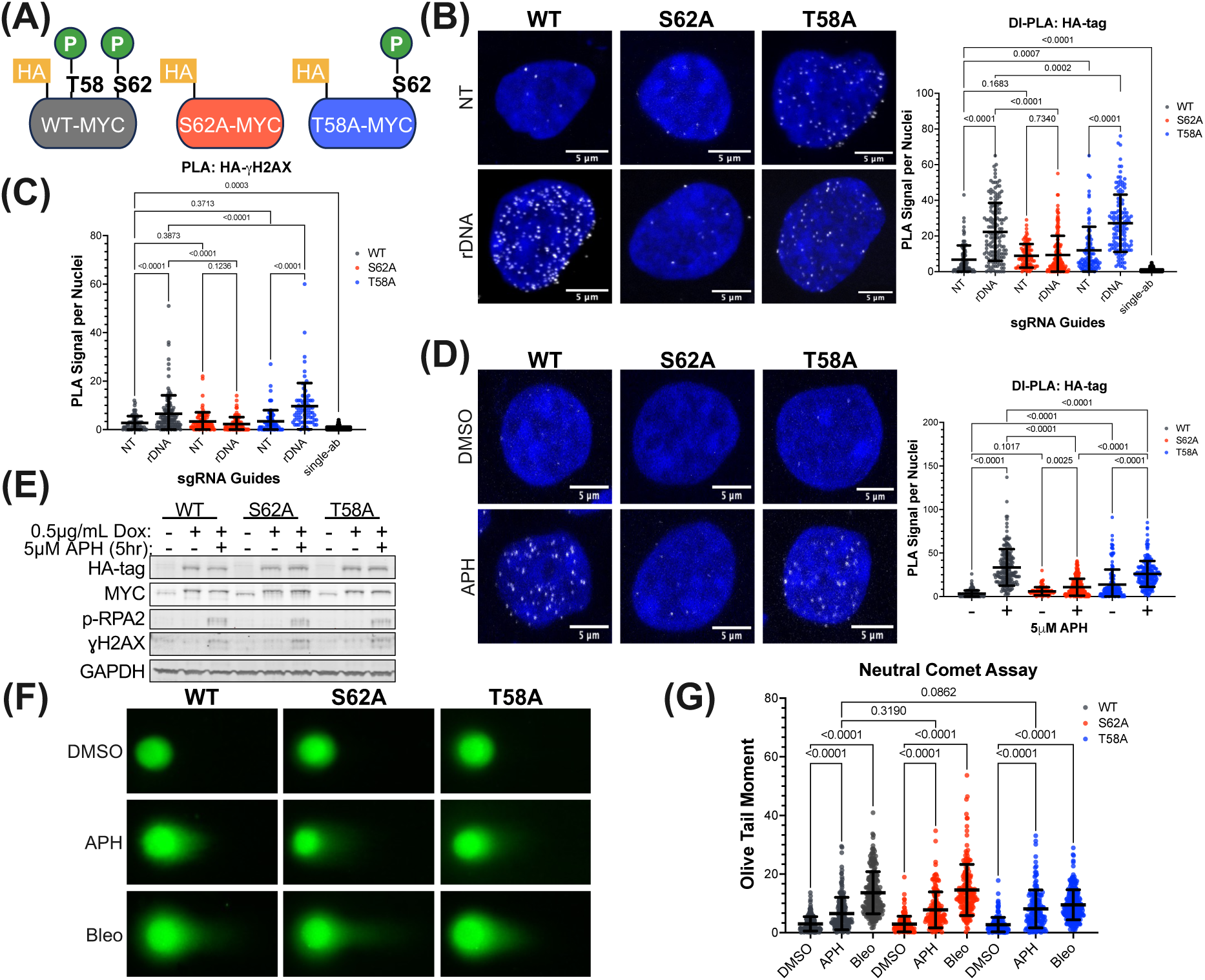
Serine 62 Phosphorylation of MYC Promotes its Association with DSBs. (A) Schema of phosphorylation status of HA-tagged WT-MYC, S62A-MYC, and T58A-MYC in doxycycline inducible HEK293 cells. (B) *Left:* Merged DI-PLA between HA-tag and biotin treated for 18-hours with 0.5μg/mL doxycycline followed by an 8-hour transient transfection with Cas9 protein and guide RNAs (sgRNA) targeting the 28S rDNA or non-targeting control (NT). PLA puncta pseudo-colored in white. *Right:* Quantification of three biological replicates of DI-PLA in. Single antibody control combines PLA counts for both primary antibodies alone treated with rDNA guides. The error bars show mean ±s.d. Statistical significance was determined by ANOVA. (C) Quantification of three biological replicates of PLA between HA-tag and γH2AX. (D) *Left*: Merged representative single-cell images of DI-PLA between HA-tag and biotin treated for 18-hours with 0.5μg/mL doxycycline followed by 5μM APH (+) or DMSO (-) for 5 hours. Right: Quantification of three biological replicates with error bars showing mean ±s.d. and statistical significance was determined by ANOVA. (E) Western Blot analysis of WT-, S62A-, and T58A-MYC in HEK293 cell lines treated with 0.5μg/mL doxycycline for 18-hours followed by 5μM APH (+) or DMSO (-) for 5 hours. (F) Neutral comet assay following the same treatment as (E). 1-hour 100 μg/mL bleomycin treatment was positive control. (G) Quantification of (F) showing the Olive Tail Moment of three biological replicates. The error bars show mean ±s.d. Statistical significance was determined by ANOVA.

Furthermore, a neutral comet assay demonstrated that the levels of DSBs induced by 5-hour APH treatment was not significantly different between cells expressing WT-, S62A-, and T58A-MYC, confirming that the number of DSBs is equivalent across the conditions (Figure 4F,G). Taken together, these data underscore the critical role of serine 62 phosphorylation in MYC’s efficient association with DSBs.

### Serine 62 Phosphorylation of MYC Regulates the Efficient Recruitment of BRCA1 and RAD51 to DSBs

To assess whether MYC could influence the recruitment of repair factors to DSBs, we initially evaluated the impact of MYC phosphorylation status on its association with BRCA1 & RAD51 as shown in Figure 2F & G. Both WT & T58A-MYC exhibited strong and comparable association with BRCA1 following APH treatment, while S62A-MYC showed a weaker induction, significantly reduced compared to WT & T58A-MYC (Figure 5A). Similarly, WT-MYC demonstrated a pronounced APH-induced increase in RAD51 association, T58A-MYC association was also significantly increased, while S62A-MYC did not show a significantly increased interaction with RAD51 (Figure 5B). Co-immunoprecipitation of flag-tagged WT-, S62A-, or T58A-MYC in APH-treated HEK293 cells confirmed the reduced interaction between MYC and RAD51 when serine 62 phosphorylation is blocked (Figure 5C). Given S62A-MYC’s diminished association with DSBs, we next examined whether serine 62 phosphorylation of MYC and its efficient recruitment to DSBs might be important for the localization of BRCA1 and RAD51 to DSBs. DI-PLA analysis of BRCA1 in WT-MYC expressing cells demonstrated a robust APH-induced increase in BRCA1 association with DSBs (Figure 5D,E). However, this induction was not observed in S62A-MYC expressing cells, with no significant difference in BRCA1 association with DSBs between DMSO and APH treatment in these cells. A similar pattern was observed with cas9-induced DSBs (Figure 5F), with the unexpected finding that BRCA1’s association with DSBs falls below NT control in S62A-MYC expressing cells, an observation that warrants further experimentation to draw a definitive conclusion. S62A-MYC expressing cells also had a marked reduction in RAD51 localization to APH-induced DSBs compared to WT-MYC expressing cells (Figure 5G,H). These results collectively demonstrate that efficient recruitment of BRCA1 and RAD51 to DSBs as well as their association with MYC is reliant on serine 62 phosphorylation of MYC.

**Figure 5:**
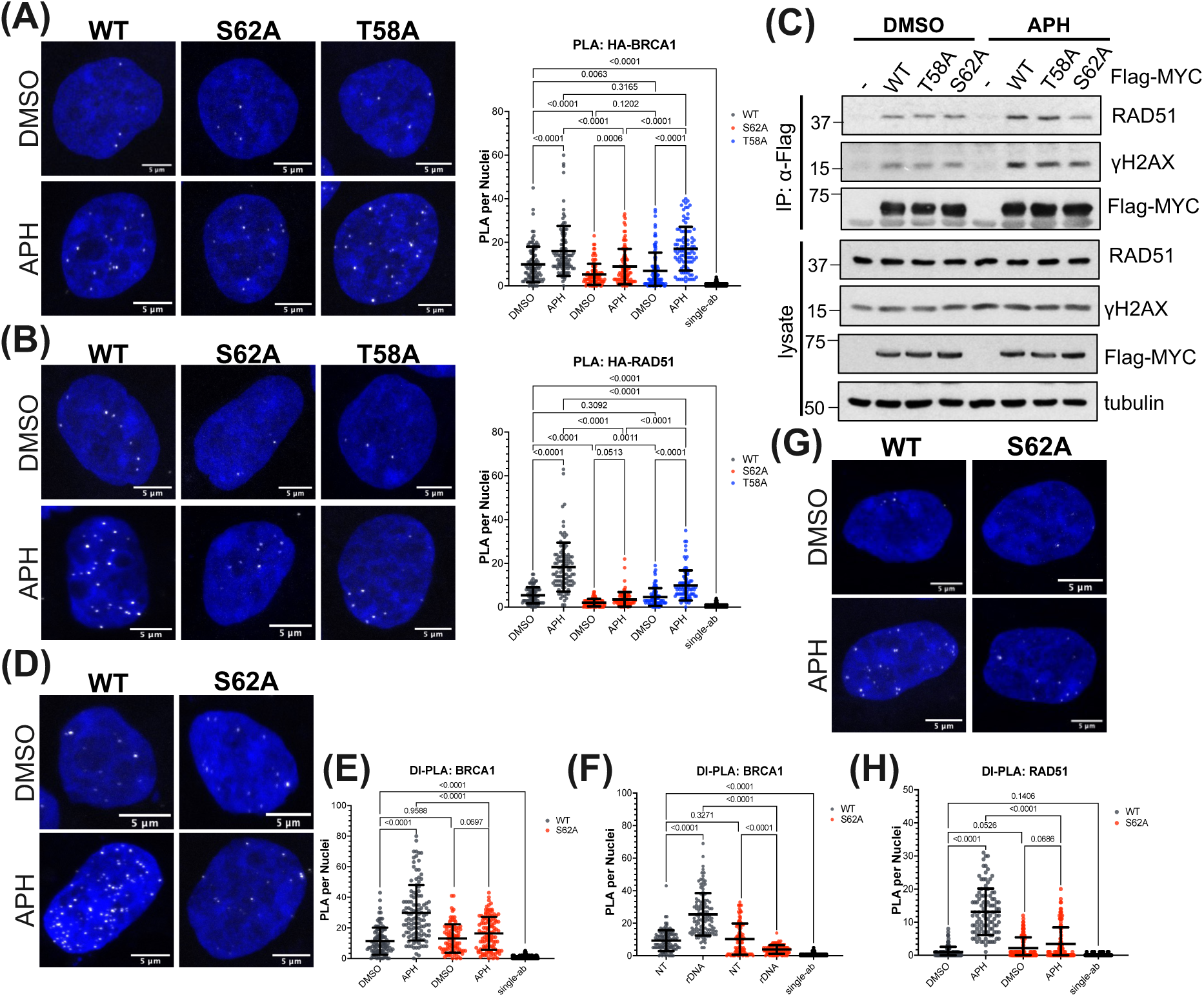
Serine 62 Phosphorylation of MYC Regulates the Efficient Recruitment of BRCA1 and RAD51 to DSBs. (A) Merged PLA between HA-tag and BRCA1 treated for 18-hours with 0.5μg/mL doxycycline followed by 5 hours of either DMSO or 5μM APH. PLA pseudo-colored in white. (B) Quantification of (A) experiment. Single antibody control combines PLA counts for both primary antibodies alone treated with 5μM APH for 5-hours. The error bars show mean ±s.d. Statistical significance was determined by ANOVA. (C) Merged PLA between HA-tag and RAD51 treated for 18-hours with 0.5μg/mL doxycycline followed by 5 hours of either DMSO or 5μM APH. PLA pseudo-colored in white. (D) quantification of (C). (E) Co-immunoprecipitation of transiently transfected Flag-tagged WT, T58A, or S62A-MYC in HEK293 cells followed by 5-hour treatment with either DMSO or 5μM APH. Anti-Flag antibody for precipitation followed by Western Blot analysis. (F) Merged DI-PLA experiment between BRCA1 and biotin in WT or S62A-MYC expressing HEK293 cells treated with 0.5μg/mL doxycycline for 18-hours followed by 5-hour treatment with either DMSO or 5μM APH. (G) Quantification of (F). (H) Merged DI-PLA experiment between RAD51 and biotin in WT or S62A-MYC expressing HEK293 cells treated with 0.5μg/mL doxycycline for 18-hours followed by 5-hour treatment with either DMSO or 5μM APH. (I) Quantification of (H).

### Phosphorylation of MYC at Serine 62 is Critical for DNA Damage Repair and Cell Survival in Response to APH-induced Stress

To investigate the biological implications of MYC’s association with DSBs and DNA repair machinery, we performed a 3-hour washout experiment following APH treatment. Consistent with Figures 4F & G showing equivalent levels of DNA damage in WT- and S62A-MYC expressing cells, APH treatment produced comparable γH2AX puncta in both WT- and S62A-MYC expressing cells (Figure 6A,B). However, while WT-MYC efficiently resolved many of those γH2AX puncta during the 3-hour washout, the number of puncta increased in S62A-MYC expressing cells. Since γH2AX is not an exclusive marker of DSBs, we performed a neutral comet assay of pre- and post-washout. In agreement, WT-MYC successfully resolved APH-induced DSBs, while S62A-MYC comet tails showed longer comet tails post-washout compared to the pre-washout (Figure 6C,D). To assess whether the deficiency in DNA damage repair observed in S62A-MYC expressing cells impacted cell survival, we conducted a 10-day colony formation assay following a 5-hour APH treatment at varying concentrations. T58A-MYC exhibited a similar number of colonies to WT-MYC, whereas S62A-MYC showed a significant reduction and an impaired ability to recover from APH-induced DNA damage (Figure 6E,F). Altogether, these findings suggest that MYC serine 62 phosphorylation impacts its association with DSBs, the recruitment of DNA repair machinery, and the promotion of DNA repair and cell survival.

**Figure 6:**
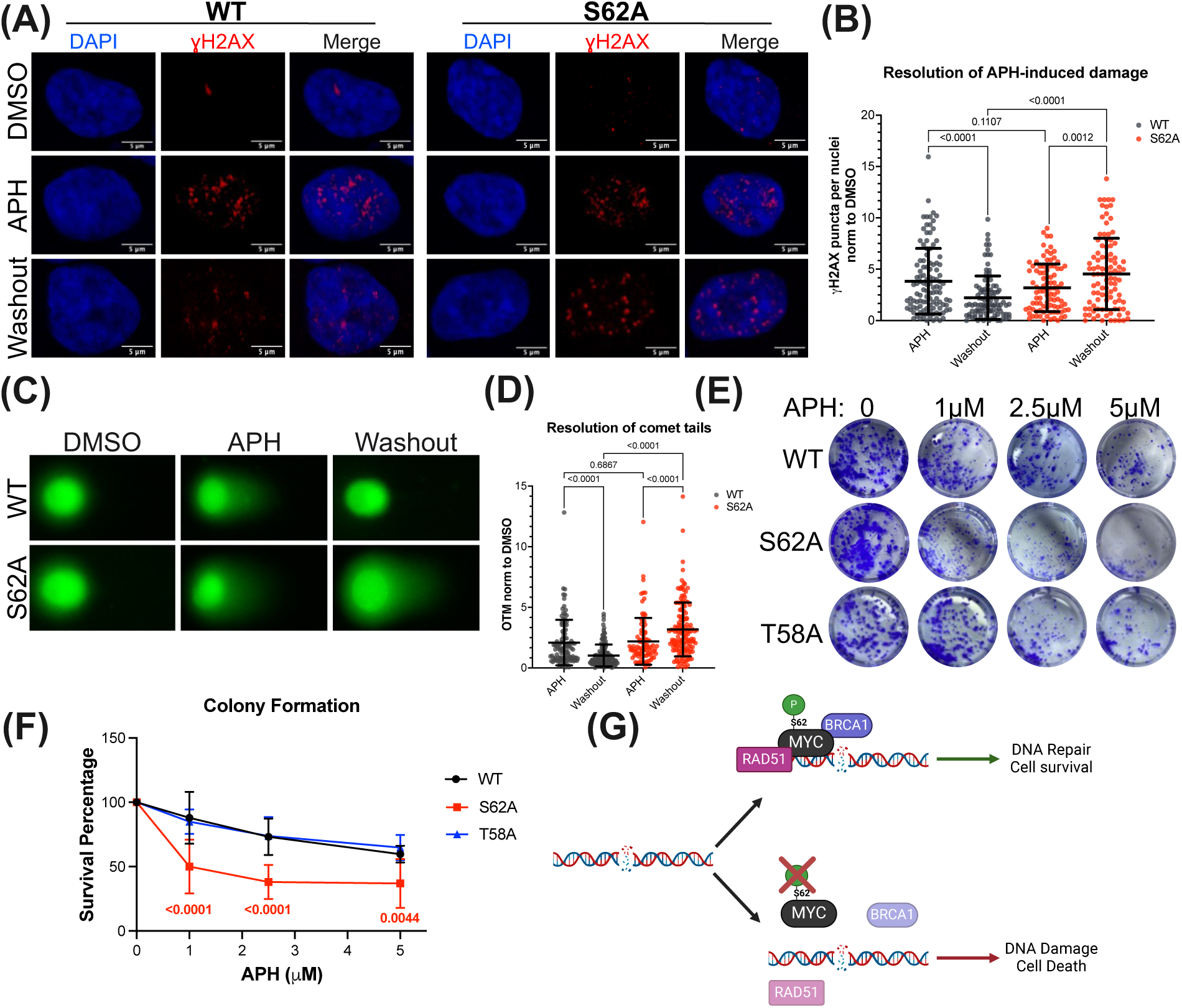
Phosphorylation of MYC at Serine 62 is Critical for DNA Damage Repair and Cell Survival in Response to APH-induced Stress. (A) Representative immunofluorescence images showing γH2AX puncta. WT- or S62A-MYC were induced by 18-hour 0.5μg/mL doxycycline treatment followed by either 5-hours of either DMSO or 5μM APH. Washout represents coverslips which were treated with 5μM APH for 5-hours followed by a media change with doxycycline for 3-hours. Statistical significance was determined by ANOVA. (B) Quantification of (A) comparing γH2AX puncta for 5-hour APH and 3-hour washout. Puncta were normalized to respective 5-hour DMSO or DMSO-washout condition. (C) Neutral comet assay of 3-hour washout post 5-hour DMSO or 5μM APH treatment in WT- or S62A-MYC HEK293 cells. (D) Quantification of (C) where washout APH treatment was normalized to washout DMSO. Statistical significance was determined by ANOVA. (E) 10-day Colony formation assay following 5-hour DMSO or APH in increasing concentrations. (F) Quantification of (E) showing the survival percentage of WT-, S62A-, and T58A-MYC expressing cells following 5-hour APH treatment. Statistical significance was determined by ANOVA. (G) Graphical summary of findings.

## Discussion

In this study, we explored the non-canonical role of MYC in DDR and its association with DSBs to elucidate new mechanisms through which MYC drives oncogenesis and cancer cell survival. Our findings underscore a role for MYC in maintaining genomic integrity through mechanisms involving a direct role in DNA repair, and we reveal that MYC’s phosphorylation at serine 62 is critical for its proximity to DSBs, recruitment of repair proteins like BRCA1 and RAD51, and overall effectiveness of DNA repair (Figure 5G).

Transcriptionally, MYC has been shown to promote genomic maintenance by regulating transcription of several DNA repair proteins including RAD50, RAD51, XRCC2, BRCA1, BRCA2, DNA-PKcs, HUS1, and Ku70 (Luoto, Meng et al. 2010). Furthermore, during S-phase, MYC upregulates expression of homologous recombination-directed repair proteins, RAD51 and HUS1 (Robson, Ward et al. 2011). In our large expression dataset from 289 human PDAC tumors, we demonstrated a positive correlation between MYC target pathway activity and both replication stress and DNA repair pathway activity (Figure 1A & B). Furthermore, high versus low MYC target pathway activity showed increased VIPER activity score for DNA repair proteins and an increase in overall survival for patients who harbor somatic DDR mutations when MYC activity is low, consistent with prior findings that heightened MYC and DDR pathways drive a subset of aggressive PDAC tumors(Link, Eng et al. 2025). This connection may not be exclusive to PDAC given that prior clinical findings have shown that MYC amplification and increased genomic instability drive breast cancer progression and aggressiveness in BRCA1-mutated tumors (Grushko, Dignam et al. 2004). The positive correlation between MYC-target pathway activity and replication stress signature, supports the idea that MYC upregulates factors to mitigate the MYC-driven stress induced by accelerated cell cycle progression (Felsher and Bishop 1999). In agreement, MYC, along with MYCN, compensates for heightened replication stress by increasing transcription of components that address stalled replication forks and facilitate DNA repair, including the MRN complexes (Chiang, Teng et al. 2003, Petroni, Sardina et al. 2016), Cohesin components (Rohban, Cerutti et al. 2017), TRIM33 (Rousseau, Einig et al. 2023), and MCM10 (Murayama, Takeuchi et al. 2021). In addition, MYC expression induces nucleotide biosynthesis genes to sustain nucleotide balance to reduce replication stress (Liu, Li et al. 2008).

Beyond MYC’s transcriptional role in genomic maintenance, emerging evidence suggests that MYC protein can directly influences DNA damage prevention and repair. In a study by Cui et al., pS62-MYC colocalized with gH2AX and DNA-PKcs/S2056 foci in irradiated HeLa cells and that MYC silencing reduced DNA repair (Cui, Fan et al. 2015). In neuroblastoma, MYCN is found at sites of heightened transcription which have a greater propensity to accumulate DNA damage. At these genomically stressed sites, the ubiquitin-specific protease USP11 was shown to preferentially bind to and stabilize de-phosphorylated threonine 58 MYCN which mediated the recruitment of BRCA1 to stalled RNAPII complexes, preventing the accumulation of deleterious R-loops (Herold, Kalb et al. 2019). In agreement, we showed that MYC’s association with DSBs and BRCA1 was greatest in T58A-MYC and diminished in S62A-MYC when compared to WT-MYC (Figures 4,5). Emphasizing that serine 62 phosphorylation is important for MYC’s association with DSBs and repair factors.

MYC elicits its cellular activity through its interactome and by recruiting and concentrating multiprotein complexes at genomic sites across the genome(Lourenco, Resetca et al. 2021, Das, Lewis et al. 2023). Our MYC-BioID2 proteomic screen detected a functional shift in MYC’s interactome under replication stress (Figure 3). Under DMSO conditions, MYC-BioID2 biotinylated more interactions with proteins involved in canonical MYC functions that drive gene expression and metabolism while APH-enriched MYC interacting proteins aligned more with DDR pathways. MYC’s interactome under cellular stress aligns with observations that MYC forms stress-induced higher-order multimeric structures around stalled replication forks, shielding them from RNAPII (Solvie, Baluapuri et al. 2022). These MYC multimers were also reported to encompass repair proteins such as FANCD2, ATR, and BRCA1, to mitigate transcription-replication conflicts and subsequent DSBs during S-phase. In addition, it was recently shown that MYCN exists in two distinct physical states depending on the phase of the cell cycle. During G1, MYCN heterodimerizes with MAX to drive transcription, whereas during S-phase, MYCN interacts with nuclear exosome targeting complexes responsible for preventing transcription-replication collisions and eliminating genotoxic RNA-structures (Papadopoulos, Solvie et al. 2022, Papadopoulos, Ha et al. 2024). These functionally distinct physical states of MYC align with numerous observations that post-translational modifications (PTMs) and protein-protein interactions partition MYC into functionally distinct “MYC-pools” which impacts its stability (Tworkowski, Salghetti et al. 2002, Guo, Li et al. 2014, Lourenco, Resetca et al. 2021). Since pS62-MYC, but not pT58-MYC, was shown to be essential for the spatial partitioning of MYC within the nucleus (Su, Pelz et al. 2018), it is plausible that phosphorylation at threonine 58 and serine 62 regulates functionally distinct MYC-pools under different cellular contexts. For example, MYC assembles and recruits a topiosome composed of topoisomerases 1 and 2 to alleviate transcription-induced topological stress (Das, Kuzin et al. 2022). However, in the presence of excessive DNA damage in cell lines, MYC is degraded and replaced with a p53-mediated topiosome, leading to proficient DDR and repair (Das, Karmakar et al. 2024). MYC also requires ubiquitination and degradation for the transfer of PAF1c to RNAPII to couple transcriptional elongation with DSB repair (Endres, Solvie et al. 2021). The notion that MYC-pools are independently regulated spatially, could help explain reports that MYC is degraded in response to DNA damage (Popov, Wanzel et al. 2007, Britton, Salles et al. 2008, Li, Challagundla et al. 2015), since a subset of more stable MYC pools, such as pS62-MYC, could allow for prolonged repair. Furthermore, since pS62-MYC is required for the efficient recruitment of BRCA1 and RAD51 to DSBs, MYC may play a role in directing homologous-directed repair. Future studies will investigate the consequences of MYC-directed repair in PDAC.

In conclusion, this study provides novel mechanistic understanding into the non-canonical role of MYC in DSB repair, demonstrating that serine 62 phosphorylation is critical for directing MYC’s efficient association with DSBs and subsequent recruitment of repair factors necessary for productive DNA repair and cell survival under stress. These insights advance our understanding of MYC’s function beyond transactional regulation, highlighting additional contributions to MYC-driven oncogenesis and resistance to DNA damaging chemotherapy.

## Methods

### Cell Lines

MIA PaCa2, U2OS, HeLa, HEK293 and MYC-mutant HA-tag HEK293TR cells were maintained in DMEM supplemented with 10% characterized fetal bovine serum (FBS), 2mM L-glutamine, and 1X penicillin/streptomycin at 37°C and 5% CO_2_. Patient derived cell line ST-00024058 was generated as previously described (Queitsch, Moore et al. 2023).

### Generation of stable inducible 293TR-MYC cells

293TR-MYC inducible cells were generated using a technique as previously described (Farrell, Pelz et al. 2013). Briefly, HEK293 cells were infected with lentivirus encoding the Tet repressor, pLenti6/TR (Invitrogen) for 12 hours. Stable clones were maintained at 5µg/mL blasticidin (Invitrogen). Clones were then infected with lentivirus expressing HA-MYC (pLenti4/TO/CMV-HA-MYC). Cells were selected with 200 µg/mL Zeocin (Invitrogen) for 10 days until clones grew out. Clones were screened for HA-MYC when treated with 1µg/mL doxycycline for 24 hours. Stable 293TR-MYC^T58A^ and 293TR-MYC^S62A^ cells were similarly constructed except with pLenti4/TO/CMV-HA-MYC^T58A^ or pLenti4/TO/CMV-HA-MYC^S62A^ respectively.

### Tissue acquisition and patient consent

Patient blood, tissues, and data were acquired with inform consent aligned with the Declaration of Helsinki and were obtained through the Oregon Pancreas Tissue Registry under Oregon Health & Science University IRB protocol #3609.

### RNA-sequencing of patient PDAC

Detailed methods for RNA preparation and sequencing can be found in Link et al (Link, Pelz et al. 2022). OHSU supplied formalin-fixed paraffin-embedded tissue sections to Tempus as part of a contract agreement. Tempus performed whole-transcriptome RNA sequencing as previously described (Beaubier, Tell et al. 2019).

### Gene Set Variation Analysis (GSVA)

GSVA analysis(Hänzelmann, Castelo et al. 2013) along with the MSigDB database v7.5.1 Hallmark gene set collection (Liberzon, Birger et al. 2015) was used to calculate Hallmark scores for all primary tumors. The replication stress gene set was derived from replication stress-induced gene expression patterns observed in Dreyer et al (Dreyer, Upstill-Goddard et al. 2021). We compiled this composite gene set from all genes the appeared in 3 or more of the 21 gene sets identified by Dreyer et al as being activated in replication stress and DNA damage response. Pearson’s correlation coefficient and p-values were calculated using the cor.test() function in R.

### VIPER analysis of high vs low Hallmark MYC-V1 score

Transcriptional regulon enrichment was analyzed using VIPER alongside the ARACNe-inferred TCGA PAAD network(Alvarez, Shen et al. 2016, Lachmann, Giorgi et al. 2016). Before running VIPER, gene expression data were normalized by median centering and scaling, and the resulting regulon scores from all primary samples were used for cohort comparisons. For the Gene Ontology (GO)(Ashburner, Ball et al. 2000, Aleksander, Balhoff et al. 2023) enrichment analysis of regulons elevated for the high Hallmark MYC-V1 target pathway cohort, we utilized the R package ClusterProfiler (v.4.6.2)(Wu, Hu et al. 2021). Primary tumors were ranked based on the HALLMARK_MYC_TARGETS_V1 GSVA score to produce quartiles. We then compared the top quartile to the bottom quartile and calculated multiple test corrected p-values (q-values) and difference in means between these quartiles for all Viper features. The enrichGO function was configured to assess GO biological process terms, with both p-value and q-value thresholds set to 0.05, and all regulons used as the background. Jaccard similarity was computed using the default setting of the pairwise_termsim.

### Cyclic Immunofluorescence multiplex imaging analysis

A PDAC tissue microarray (TMA) was created at OHSU using FFPE blocks from tumors with 1-2 cores per tumor from 34 primary tumors, totaling 55 cores. Immunofluorescence preparation and analysis conducted as previously described(Eng, Bucher et al. 2022). Briefly, images were scanned with Zeiss Axioscan Z1, acquired, stitched, and exported as TIFF format with Zeiss Zen Blue software (v.2.3). Image registration was performed using MATLAB (v.9.11.0), and cellular segmentation was carried out using either Cellpose(Stringer, Wang et al. 2021) or Mesmer(Greenwald, Miller et al. 2022). Unsupervised clustering of individual cell mean fluorescence intensity was used to classify cell types, using the Leiden algorithm implemented in scanpy (v.1.9.3)(Wolf, Angerer et al. 2018). Pearson correlation r of tumoral single-cell mean intensity of DNA damage and proliferation markers with pMYC was calculated in each core. For hypothesis testing, Pearson’s r values were transformed with Fisher’s Z and 95% confidence intervals calculated.

### DNA Damage In Situ Ligation followed Proximity Ligation Assay (DI-PLA)

This protocol is adapted from Galbiati et al (Galbiati and d’Adda di Fagagna 2019). Cells are grown on 13mm coverslips and fixed in 4% PFA for 10 minutes at room temperature followed by two washes with PBS.

#### DI-PLA: Blunting

Coverslips are washed twice for 5 minutes with NEB2 buffer (50mM NaCl, 10mM Tris-HCl pH 8, 10mM MgCl_2_, 1mM DTT, 0.1% Triton X-100) and twice for 5 minutes with Blunting buffer (50mM NaCl, 10mM Tris-HCl pH 7.5, 10mM MgCl_2_, 5mM DTT, 0.025% Triton X-100). Coverslips are then inverted onto a 35μL drop on parafilm of NEB Blunting Reaction (NEB, E1201): (1mM dNTPs, 1X Blunting Buffer, 0.2mg/mL BSA, 1X Blunting Enzyme). Coverslips are incubated in a dark humidity chamber for 1hr at room temperature.

#### DI-PLA: Ligation

Coverslips are washed twice for 5 minutes with NEB2 buffer (50mM NaCl, 10mM Tris-HCl pH 8, 10mM MgCl_2_, 1mM DTT, 0.1% Triton X-100) the twice for 5 minutes with ligation buffer (50mM Tris-HCl pH 7.5, 10mM MgCl_2_, 10mM DTT, 1mM ATP). Coverslips are then inverted onto a 50μL drop on parafilm of Ligation Reaction (0.1μM DI-PLA Linker, 1X T4 Ligation Buffer (NEB, B0202), 1mM ATP, 0.2 mg/mL BSA, 1X T4 Ligase (NEB, M0202)) overnight at 4°C in dark humidity chamber followed by proximity ligation assay between biotin and protein of interest.

#### DI-PLA Linker

5’-TACTACCTCGAGAGTTACGCTAGGGATAACAGGGTAATATAGTTT [BtndT] TTTCTATATTACCCTGTTATCCCTAGCGTAACTCTCGAGGTAGTA -3’

### Proximity Ligation Assay

Proximity Ligation Assay was performed without deviation from manufacturer’s instructions (DUO92008). Single-antibody controls were performed to ensure specificity of antibodies. Coverslips were washed in a 0.5mL volume and reactions were performed by inverting the coverslip onto a 35μL drop on parafilm. Following the proximity ligation reaction, cells stained with DAPI (0.2ug/mL) for 3 minutes followed by one wash in PBS and one water wash. The cells were then inverted and mounted on glass coverslips with 15μL of prolong gold mounting media (LifeTech, P36934) & were cured overnight in the dark at room temperature. A minimum of 30 cells were imaged per replicate at 63X on a Zeiss LSM880 confocal microscope and analyzed with CellProfiler. Statistical significance was performed using two-tailed student’s t-test or one-way ANOVA with multiple comparisons. Antibodies used are as follows: MYC (Abcam, 32072), pS62-MYC (Abcam, 78318), Biotin (Sigma, B7653), Biotin (Abcam, 53494), RAD51 (Abcam, 133534), HA-tag (ABM, G036), pRPA2 (Novus, NB100-544), γH2AX (Invitrogen, MA12022), γH2AX (Cell Signaling, 9718S), BRCA1 (Santa Cruz, 6954), BRCA1 (Sigma, 07-434-MI).

### Cas9-Transfection

Cells were cultured on glass coverslips and transiently transfected using Lipofectamine CRISPRMAX (Thermo, #CMAX00015) with TrueCut Cas9 Protein v2 (Thermo, #A36499) and synthetic guide RNA from Invitrogen TrueGuide Synthetic sgRNA (Cat#: 35514) or Negative Control (Cat#: A35526) following manufacturer’s instructions. After 8-hour incubation, cells were fixed in 4% paraformaldehyde for 10 minutes at room temperature, washed three times with PBS, and stored at 4°C. The following two rDNA guides were used in a 1:1 ratio of rDNA guide 1: CGAGAGAACAGCAGGCCCGC and rDNA guide 3: GATTTCCAGGGACGGCGCCT.

### Western Blot

MYC-Mutant expressing HEK293 cells were seeded into 6-well chamber well. The next day, cells were treated with 0.5μg/mL doxycycline for 18 hours then treated with 5μM aphidicolin for 5 hours. Cells were then washed three times with DPBS and flash frozen at -70°C. Cells were then thawed and scraped in lysis buffer (20mM Tris-HCl pH7.5, 50mM NaCl, 0.5% Triton X-100, 0.5% DOC, 0.5% SDS, 1mM EDTA, protease inhibitor (Millipore Sigma: 5892791001), and phosphatase inhibitor (Millipore Sigma: 4906837001). Lysates were then sonicated with a Branson Sonifier 450 for 10 pulses, Duty factor of 20, and an output of 2. Protein content was then quantified and 25μg of protein was boiled in 1X SDS buffer (50mM Tris-Cl pH 6.8, 2%SDS, 6% glycerol, 5% 2-mercaptoethanol). Samples were loaded into a 4-12% Bis-Tris Citerion Gel (BioRad: 345-0123) and run for 1.5 hours at 180V on ice in XT-MOPS (BioRad: 161-0788). The gel is then transferred onto an Immobilin PVDF membrane (Fisher Scientific: IPFL00010) for 90 minutes at 400mA. Membrane is then blocked for 1 hour at room temperature in Aquablock (Arlington Scientific: NC2580736) then incubated overnight with primary antibody. The following day, the membrane is washed for three times with TBST and incubated with secondary Licor antibody for 1 hour in the dark room temperature and imaged on a Licor Odyssey scanner. Antibodies used are as follows: MYC (Abcam, 32072), HA-tag (ABM, G036), pRPA2 (Novus, NB100-544), γH2AX (Invitrogen, MA12022), GAPDH (Fisher, AM4300). Anti-rabbit IgG 800CW (VMR, 102673-330), Anti-mouse IgG 680RD (LI-COR, 926-68072), Anti-mouse IgG 800CW (Fisher, NC9401841).

### BioID2 Cloning

BioID2 sequence with a five-glycine linker (Addgene: 74224) was cloned at the N-terminus of C-MYC (Addgene: 16011) and inserted into pcDNA4/TO (Invitrogen: V102020) via HindIII (NEB: R0104T) and NotI (NEB: R0189L). The HA-BioID2 only control sequence (Invitrogen: 74224) was cloned into pcDNA4/TO via HindIII and BspeI (NEB: R0540) Each plasmid was independently transfected using Lipofectamine 3000 (Thermo Fisher Scientific L3000-015) into HEK293 cells which contained pcDNA6/TR (Invitrogen: V102520) under blasticidin selection at 20ug/mL. Single cell clones were isolated after several days of Zeocin selection at 200ug/mL. MYC-BioID2 positive clones were identified by western blot for biotin and MYC expression upon doxycycline induction (1ug/mL for 24 hours) with Strepavidin 680 (LI-COR Biosciences 926-68079) and Y69 (Abcam ab32072). HA-BioID2 positive clones were screened using a HA-tag antibody (Applied Biological Materials G036).

### MYC-BioID2 Assay

Cells were plated on two 15-cm dishes per condition to achieve 80% confluency upon treatment. Cells were treated for 24-hours with 1 µg/mL doxycycline in combination with either 5 µM APH or DMSO vehicle control. For biotinylation, cells were treated with 50 µM biotin (Millipore Sigma: B4501) for 18 hours. After treatment, cells were washed twice with PBS and frozen at -70°C and stored until three biological replicates were obtained. Cells were then lysed with 1.5 mL/dish of lysis buffer (50 mM Tris-Cl, pH 7.4, 150 mM NaCl, 1% Triton X-100, 0.1% SDS; 1 mM DTT, protease inhibitor (Millipore Sigma: 5892791001), and phosphatase inhibitor (Millipore Sigma: 4906837001), and incubated on ice for 10 minutes. Lysates were scraped, pooled, and sonicated with a Branson Sonifier 450 for two 30-pulse sessions at a 30% duty cycle and 1.5 output, with a 2-minute ice interval. Samples were diluted with 2.6 mL pre-chilled 50 mM Tris-Cl (pH 7.4) and sonicated for an additional 30 pulses. Lysates were centrifuged at 16,500 x g for 10 minutes at 4°C to remove debris. 150 µL/condition of Streptavidin-agarose beads (Millipore Sigma: 69203-3) were equilibrated in a 1:1 mixture of lysis buffer and 50 mM Tris-Cl (pH 7.4) and briefly spun down at 8,000 rpm for 2 minutes. Supernatants were transferred to the beads and incubated overnight on a rotator at 4°C. After incubation, beads were washed twice with wash buffer 1 (50 mM HEPES pH 7.5, 400 mM NaCl, 1% Triton X-100, 0.1% deoxycholic acid, 1 mM EDTA) for eight minutes, twice with wash buffer 2 (10 mM Tris-Cl pH 7.4, 500 mM LiCl, 0.5% NP-40, 0.5% deoxycholic acid, 1 mM EDTA) at room temperature. Beads were washed once in 50 mM Tris-Cl (pH 7.4) followed by three washes with 100mM ammonium bicarbonate. Beads were resuspended in 100 µL of 100mM ammonium bicarbonate (pH 8) and underwent trypsinization with the addition of 15µL of 80ng/µL trypsin (1.6ug) in 50mM Triethyl ammonium bicarbonate and incubated for 17 hours at 37°C with shaking. After trypsinization, beads were isolated, supernatant removed and filtered with 0.22µm Millipore filter. Filtered sample was dried and dissolved in 20µL of 5% formic acid and injected into Thermo QExactive HF mass spectrometer and run with the 90min LC/MS method. Survey mass spectra were acquired over m/z 375−1400 at 120,000 resolution (m/z 200) and data-dependent acquisition selected the top 10 most abundant precursor ions for tandem mass spectrometry by HCD fragmentation using an isolation width of 1.2 m/z, normalized collision energy of 30, and a resolution of 30,000. Dynamic exclusion was set to auto, charge state for MS/MS +2 to +7, maximum ion time 100ms, minimum AGC target of 3 x 106 in MS1 mode and 5 x 103 in MS2 mode. Mass spectrometry data from all samples was processed using COMET/PAWS against Uniprot Human database. Comet (v. 2016.01, rev. 3)(Eng, Jahan et al. 2013) was used to search MS2 Spectra against a January 2024 version of canonical FASTA protein database containing human uniprot sequences, and concatenated sequence-reversed entries to estimate error thresholds. Comet searches for all samples were performed with trypsin enzyme specificity with monoisotopic parent ion mass tolerance set to 1.25 Da and monoisotopic fragment ion mass tolerance set at 1.0005 Da and a variable modification of +15.9949 Da on Methionine residues.

### Protein interaction ontology analysis

BioID2-only detected spectral counts were subtracted from MYC-BioID2 to remove background biotinylation and spectral counts from MYC-BioID2 samples treated with either DMSO or APH were normalized by scaling the average total spectral counts across samples. Log2 fold changes were computed between APH- and DMSO-treated samples, and statistical significance was assessed using two-sided t-test followed by a Benjamini-Hochberg correction. Ontological analysis of targets which pass the 2-fold differential abundance threshold was determined with the BinGO tool in Cytoscape(Shannon, Markiel et al. 2003, Maere, Heymans et al. 2005, Doncheva, Morris et al. 2019).

### Neutral comet assay

Cells were plated in 6-well plate and treated with either DMSO control or 5mM aphidicolin for 5-hours. A positive control of 1-hour 100mg/mL bleomycin was included. Glass slides were pre-coated in 1% normal melting point agarose and dried at room temperature. Cells were trypsinized, counted, and resuspended in PBS to 0.35 × 10^6^ cells/mL on ice. Cell suspensions were combined with molten 1% low-melting-point agarose at a 1:10 (v/v) ratio, then 200 µL of this mixture was applied to pre-coated slides labeled in pencil. Coverslips were added for even distribution, and slides were solidified at 4 °C for 10 minutes in the dark. Slides were then submerged in 4°C lysis buffer (2.5 M NaCl, 100 mM EDTA, 10 mM Tris, 200 mM NaOH, 1% sarcosinate, and 1% Triton X-100, pH 10) for 18-hours. Slides were equilibrated in neutral electrophoresis buffer (100mM Tris and 300mM sodium acetate, pH 9) at 4°C for 30 minutes and transferred to an electrophoresis chamber with chilled buffer.

Electrophoresis was conducted at 1 V/cm for 30 minutes at 4 °C. Slides were immersed in DNA precipitation solution (1M ammonium acetate in 80% ethanol) for 30 minutes, followed by a 30-minute rinse in 70% ethanol at room temperature. Slides were dried at 37 °C, then stained with SYBR Green (1:2000, Thermo cat#: S33102) for 15 minutes in the dark. Following a brief rinse in dH2O, slides were dried partially and stored. Images were acquired at 10x using a BioTek Cytation 5 microscope. Comets analyzed using ImageJ software plugin, OpenComet(Gyori, Venkatachalam et al. 2014). Olive Tail Moment (OTM) was calculated using the following equation.

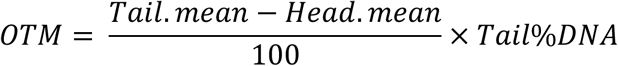

### Co-Immunoprecipitation

HEK293 cells were transfected with plasmids using Lipofectamine 2000 (Life Technologies) following the manufacturers’ protocols. The cells were harvested at 36– 48 h post-transfection, washed with PBS and then lysed in ice-cold lysis buffer consisting of 30 mM Tris-HCl (pH 8.0), 0.5% Nonidet P-40, 1 mM EDTA, 1 mM EGTA, 200 mM NaCl, 1 mM phenylmethylsulfonyl fluoride, 1 mM DTT, 1 μg/mL pepstatin A, and 1 mM leupeptin and 1mM β-Glycerophosphate with sonication. Equal amounts of clear cell lysate were incubated with anti-Flag M2 agarose beads for 5 hours at 4°C. The beads were washed 5 times with lysis buffer. Bound proteins were detected by immunoblot using antibodies, as indicated in Figure legends.

### Immunofluorescence staining

HEK293 cells with either HA-MYC or S62A-MYC were plated on poly-D-lysine treated glass coverslips and incubated for 24-hours. Expression was induced with 0.5µg/mL doxycycline for 18-hours then treated with 5µM APH or DMSO for 5-hours. Cells were then either fixed with 4% paraformaldehyde for 10 minutes or media change with dox for 3-hours followed by fixation. Coverslips were then permeabilized with 0.25% Triton X-100 for 10 minutes followed by a 1-hour block (10% goat serum, 0.1% Triton X-100). Coverslips were then incubated in primary antibody (gH2AX, Cell Signaling, 9718S) overnight at RT. Cells were then washed in three times in 1X PBS for 10 minutes and incubated in secondary antibody (Jackson Immuno, 111-565-144) for 2 hours at RT. Coverslips are washed again for three times in 1X PBS and stained with 0.2µg/mL DAPI for 3 minutes, mounted (LifeTech, P36934), and cured overnight in the dark at room temperature. Coverslips were imaged at 63X on a Zeiss LSM880 confocal microscope and puncta were analyzed with CellProfiler (www.cellprofiler.org) (Stirling, Swain-Bowden et al. 2021).

### Colony formation assay

Cells were treated in a 6-well plate then trypsinized, counted, and plated at 1,000 cells/well of a 12-well plate. Media was changed every three days with the addition of 0.5µg/mL doxycycline. After 10 days, the media was removed and stained with 0.5% crystal violet in 50% ethanol for 30 minutes. The excess dye was then gently washed away with slow-running water. The plates were then dried at RT and colonies were counted.

## Acknowledgements

The authors acknowledge expert technical assistance by the OHSU Advanced Light Microscopy Core (RRID:SCR_009961), supported by the OHSU Knight Cancer Institute (NIH P30 CA069533). In addition, we’d like to thank OHSU’s shared proteomics core for support and guidance on this project. The authors would also like to acknowledge and thank Drs. Moriah R. Arnold and Vivek K. Unni for assistance in the Cas9-induced DNA damage experiments. This manuscript was financially supported by the following: NCI U01CA224012 (RCS & JRB), U01CA278923 and R01s CA186241, CA196228 and DoD PA210068, R01CA287672 (JRB), R01 CA212600 (JRB), R21 CA263996 (RCS & JRB); the Brenden-Colson Center for Pancreatic Care and the Krista L. Lake Endowed Chair. GMC was supported, in part, by the Knight Cancer Institute stipend award.

## Author Contributions

GMC and RCS contributed to the project design; GMC performed and analyzed all experiments unless otherwise stated; CP analyzed RNA-sequencing data; KC stained tumor microarray and JE analyzed tumor microarray; CDL is the principal investigator of the clinical trial which lead to the created of PDAC patient-derived cell lines; JRB developed technology for deriving patient-derived cell lines; AS assisted in the generation and classification of the patient-derived PDAC cell line, ST-00024058; CJD & GMC designed and executed MYC-BioID2 experiments and analysis with bioinformatic help from CP; XXS & MD performed Co-Immunoprecipitation experiment. GMC wrote the paper with edits and revisions from RCS and CJD. All authors have approved the current version of the manuscript.

